# Characterisation of two *Toxoplasma* PROPPINs homologous to Atg18/WIPI suggests they have evolved distinct specialised functions

**DOI:** 10.1101/236067

**Authors:** Hoa Mai Nguyen, Shuxian Liu, Wassim Daher, Feng Tan, Sébastien Besteiro

## Abstract

*Toxoplasma gondii* is a parasitic protist possessing a limited set of proteins involved in the autophagy pathway, a self-degradative machinery for protein and organelle recycling. This distant eukaryote has even repurposed part of this machinery, centered on protein ATG8, for a non-degradative function related to the maintenance of the apicoplast, a parasite-specific organelle. However, some evidence also suggest *Toxoplasma* is able to generate autophagic vesicles upon stress, and that some autophagy-related proteins, such as ATG9, might be involved solely in the canonical autophagy function. Here, we have characterised two *Toxoplasma* proteins containing WD-40 repeat that can bind lipids for their recruitment to vesicular structures upon stress. They belong to the PROPPIN family and are homologues to ATG18/WIPI, which are known to be important for the autophagic process. We conducted a functional analysis of these two *Toxoplasma* PROPPINs. One of them is dispensable for normal *in vitro* growth, although it may play a role for parasite survival in specific stress conditions or for parasite fitness in the host, through a canonical autophagy-related function. The other, however, seems important for parasite viability in normal growth conditions and could be primarily involved in a non-canonical function. These divergent roles for two proteins from the same family illustrate the functional versatility of the autophagy-related machinery in *Toxoplasma.*

## INTRODUCTION

Macroautophagy (simply referred to as autophagy thereafter), is a life-promoting lysosomal degradation pathway required for maintaining cellular homeostasis and surviving external stresses such as periods of nutrient deprivation [1], This process allows the degradation and recycling of cellular components through their segregation into multi-membrane vesicles called autophagosomes, which will eventually fuse with lysosomes. The understanding of the molecular processes underlying autophagy was revolutionised by the discovery of so-called Atg (AuTophaGy- related) proteins in yeast [2,3], many of which are conserved in humans and other eukaryotes.

A subset of the Atg proteins constitutes the core molecular machinery required for autophagosome formation, which is a highly regulated process. Early steps involve a positive regulation by the class III phosphatidylinositol 3-kinase (PtdIns3K) complex [4], responsible for the production of phosphatidylinositol-3-phosphate (PtdIns3P). This lipid is important for the correct localisation of some of the Atg proteins like Atg18, which will in turn enable the recruitment of proteins such as Atg9, and subsequently of Atg8 to the nascent autophagosome. Atg8 (also called LC3 (light-chain 3) in mammals) is a protein with structural homology to ubiquitin, which is essential for autophagosome formation [5]. Its association with the autophagosomal membranes depends on ubiquitin-like conjugations system, comprising proteins such as Atg7 (E1-like) and Atg3 (E2-like) [6]. Because it remains associated with autophagosomal membranes until their degradation in a lytic compartment, Atg8 has been widely used as a marker for identifying and quantifying of autophagosomes in eukaryotes.

Earlier markers for autophagosome formation include members of the β-propellers that bind phosphoinositides (PROPPIN) family. These proteins are part of an evolutionarily conserved family of proteins that includes members with an important PtdIns3P-dependent effector function in autophagy, like the Atg18 protein in yeast and the WIPI (for WD-repeat protein Interacting with Phosphoinositides) proteins in mammals [7], They have multiple WD-40 repeats that fold to form seven bladed β-propellers and contain a conserved motif for interaction with phospholipids [8-10]. Repeated WD-40 motifs form β-propeller structures that act as sites for protein-protein interaction, and proteins containing these motifs serve as platforms for the assembly of protein complexes or as mediators of stable or transient interactions among other proteins [11].

Three PROPPIN proteins have been identified in budding yeast: Atg18, Atg21, and Hsv2 (homologous with swollen vacuole phenotype 2). They are involved in different subtypes of autophagy. Atg18 is a core protein required for proper autophagy function and for the yeast-specific cytoplasm-to vacuole (Cvt) trafficking pathway [12]. Atg18 also associates with the vacuole (yeast’s lytic compartment) in a PtdIns(3,5)P_2_-dependent way, where it is involved in non-autophagic functions like retrograde vesicular transport from the vacuole to the Golgi [13]. Atg21 is essentially involved in the Cvt pathway [14], although it should be noted both Atg18 and Atg21 have a similar ability to recruit Atg9 to the nascent autophagosomal membrane [15]. Hsv2 is involved in piecemeal microautophagy of the nucleus, where non-essential parts of the nucleus are removed by autophagy [12]. Mammals have four PROPPIN proteins, namely WIPI1–4. This variety of PROPPINs (with an increased complexity, including splice variants) appear to have non-redundant, yet important, functional contributions to the autophagy process [7], The individual contributions of the WIPI members to the process at the nascent autophagosome are, however, not completely elucidated.

*Toxoplasma gondii* is a widespread obligate intracellular parasitic protist that is essentially harmless to immunocompetent individuals, although developing fetuses and immunocompromised individuals can develop severe, life-threatening, infections [16]. The rapidly replicating and disease-causing forms of the parasite are called tachyzoites. As an early-diverging eukaryote, it is perhaps not very surprising to notice *Toxoplasma* contains a limited repertoire of autophagy-related proteins compared with yeast and mammals [17]. Tachyzoites have nevertheless been shown to be able to generate autophagosomes in response to nutrient deprivation [18,19] and stress of the endoplasmic reticulum [20], suggesting the presence of a functional catabolic autophagy pathway. Quite surprisingly TgATG8, *Toxoplasma* Atg8 homologue, also localises to an apicomplexan-specific plastid called the apicoplast, where it plays an important function in organelle inheritance during parasite division [21]. Other proteins regulating TgATG8 membrane association, such as TgATG3 and TgATG4, have also been shown to be important for apicoplast homeostasis [18,22]: it thus seems *Toxoplasma* has been repurposing part of the autophagy-related machinery for a non-canonical function related to its plastid [23]. On the other hand, the *Toxoplasma* homologue of Atg9, a protein involved in the early steps of autophagosome biogenesis, is not needed for proper apicoplast function, but important for sustaining stress conditions instead [24], Overall, this suggests a dual involvement of TgATG8 and its associated membrane conjugation machinery to canonical degradative autophagy, as well as a non-canonical apicoplast related function, while early players in autophagosome formation might be involved only in canonical autophagy [25].

To verify this, we sought to investigate the function of Atg18 homologues in the parasite. Indeed, in other eukaryotes both Atg18 and Atg9 are known to localise to nascent autophagosomes and to be required for their expansion [26]. Moreover, the localisation of Atg18 to nascent autophagosomes depends on the presence of PtdIns3P [27], a lipid which is also known to be important for apicoplast biogenesis in *Toxoplasma* [28]. It was thus important to assess a possible involvement of *Toxoplasma* Atg18 homologues in canonical autophagy or an apicoplast-related function.

## RESULTS

### The *Toxoplasma gondii* genome codes for two WD-40 repeat proteins of the PROPPIN family

We performed homology searches in the *T. gondii* genomic database (http://www.toxodb.org) using Atg18 from *Saccharomyces cerevisiae* as a query, and we identified two putative homologues: TGGT1_288600 and TGGT1_220160 (Fig 1A). Reverse BLAST analysis showed these two proteins had homology for members of the yeast PROPPIN family, namely yeast Hsv2 and Atg18. Both *Toxoplasma* proteins contain a putative WD-40 repeat domain (Fig 1B) within which the conserved FRRG lipid-binding motif of PROPPINs [8] was identified (Fig 1B). A phylogenetic analysis of the WD-40 repeat domain showed unambiguously that the two *Toxoplasma* homologues belonged to the PROPPINs family (the WD-40 domain of known seven-bladed β-propeller yeast protein Tup1p [29] was included in the analysis as an outlier). TgPROP1 and TgPROP2 segregated together with the Atg18/WIPIl- WIPI2 group, although it was difficult to unambiguously identify their respective homologues in the PROPPIN family. Consequently, we thus named them TgPROP1 (TGGT1_288600) and TgPROP2 (TGGT1_220160).

**Fig 1.**
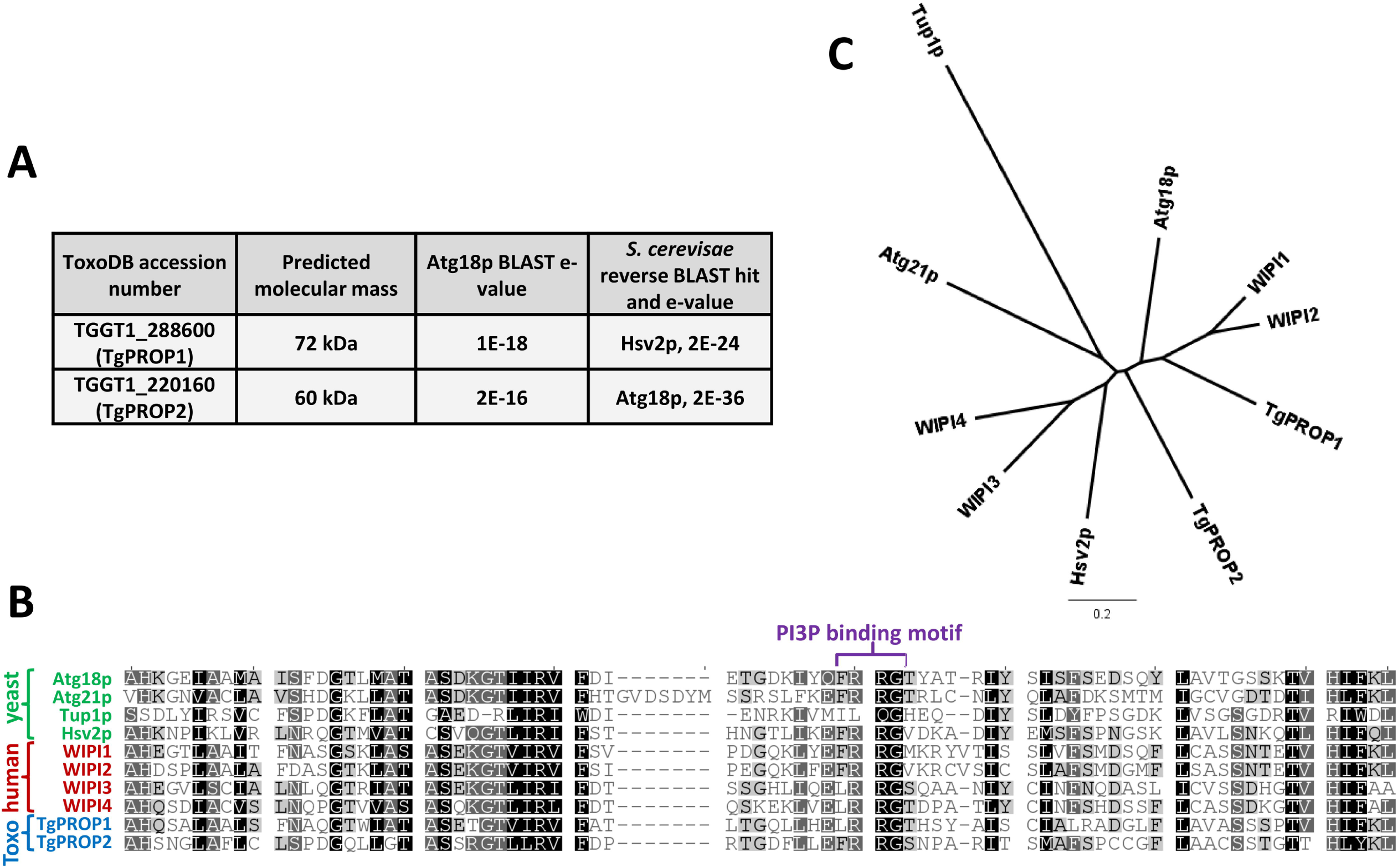
*T. gondii* contains two WD-40 repeat proteins with homology to Atg18 that belong to the PROPPIN family. **A)** Basic Local Alignment Search Tool (BLAST) search in the *T. gondii* genomic database (http://www.toxoDB.org) using *S. cerevisiae* Atg18p as a query retrieved two potential candidates we named TgPROP1 and TgPROP2. **B)** Alignment of the core WD-40 repeat region from TgPROP1 and TgPROP2 with that of yeast and human PROPPINs. Another yeast WD-40 repeats-containing protein, Tup1p, was also included in the analysis. Amino acids sequences were aligned using the Multiple Sequence Comparison by Log-Expectation (MUSCLE) algorithm. **C)** Unrooted phylogenetic tree, built from the alignment described in **B),** using unweighted pair group method with arithmetic mean. WD-40 repeats-containing protein Tup1p, serves as an outlier in the analysis. Scale bar represents 0.2 residue substitution per site.

### TgPROP1 and TgPROP2 are cytoplasmic proteins that associate with vesicles upon nutrient deprivation

To investigate the localisation of TgPROP1 and TgPROP2, we cloned their respective cDNA in fusion with the sequence coding for the dimeric Tomato (dT) red fluorescent protein. We isolated stable cell lines expressing, in addition to their native copies, either C-terminally tagged TgPROP1 or TgPROP2 (named TgPROP1-dT and TgPROP2-dT, respectively). By immunoblot (Fig 2A), we detected main products corresponding to the dT-tagged proteins, with an apparent molecular mass larger than the predicted one (predicted masses for the dT fusions are: 114 kDa for TgPROP1 and 102 kDa for TgPROP2). Fluorescence microscopy analysis showed both *Toxoplasma* PROPPINs localise to the cytoplasm (Fig 2B). As mentioned earlier, both proteins can potentially bind PtdIns3P, a lipid thought to be relatively abundant at the apicoplast membrane [28]; however co-labeling with apicoplast marker TgATRX1 [30] revealed no particular enrichment of the TgPROP1 or TgPROP2 signals at the organelle (Fig 2B).

**Fig 2.**
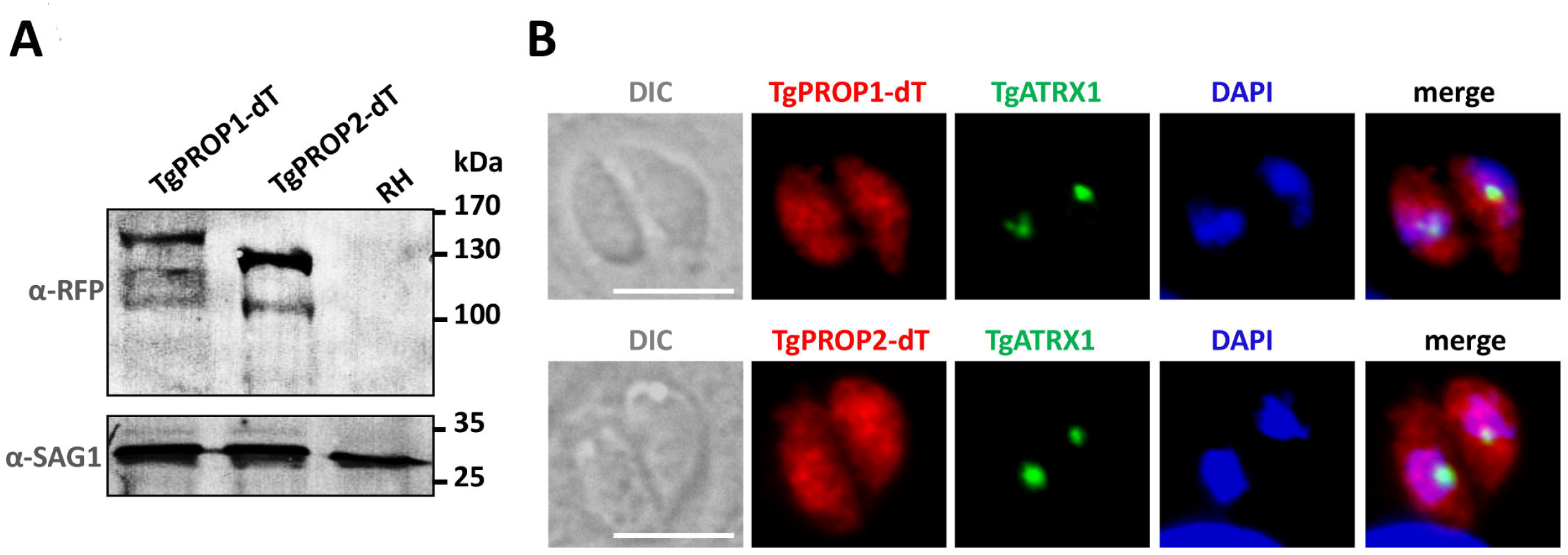
C-terminal tagging of TgPROP1 and TgPROP2 with the tandem dimer Tomato fluorescent protein. **A)** Immunoblot analysis using anti-RFP antibody to detect dT-tagged TgPROP1 and TgPROP2 in protein extracts from transgenic cell lines or the parental RH cell line. Anti-SAG1 was used as a loading control. **B)** Localisation of tandem dimer Tomato-tagged TgPROP1 and TgPROP2 in intracellular parasites by fluorescence microscopy. Co-staining was made with apicoplast marker TgATRX1. DNA was stained with DAPI. Scale bar= 5 μm.

In order to investigate a possible recruitment of TgPROP1 or TgPROP2 on autophagosomal structures, we induced a nutrient stress by incubating extracellular tachyzoites in a medium with glucose, but without amino acid. This was used before to induce the biogenesis of TgATG8-decorated autophagosomes in the parasites [18]. In these conditions, a proportion of the parasites expressing TgPROP1-dT or TgPROP2-dT displayed a puncta-like signal in addition to the cytoplasmic localisation (Fig 3A). Again, using TgATRX1 labeling, we could show this signal is in the vicinity of, but distinct from, the apicoplast (Fig 3A). To assess if these puncta could correspond to autophagic vesicles, we generated cell lines co-expressing GFP-TgATG8 together with TgPROP1-dT or TgPROP2-dT. In starvation conditions, these parasites displayed TgPROP puncta that were generally co-labeled with TgATG8, although not all TgATG8 puncta were labeled with TgPROP1 or TgPROP2 (Fig 3B). We next performed a time course experiment to quantify TgPROP1/TgPROP2- or TgATG8- positive puncta over time spent in starvation conditions. First, the percentage of parasites harboring TgPROP1- or TgPROP2-positive puncta increased with time spent in starvation conditions, likewiseTgATG8- positive puncta (Fig 3C), hinting they could be of autophagosomal nature. Second, less parasites were displaying TgPROP2-positive puncta (and even less TgPROP1-positive puncta) than TgATG8-positive puncta, which is in accordance with our previous observation that not all TgATG8-positive puncta are co-labelled with TgPROP1 or TgPROP2 (Fig 3B), suggesting TgPROP1/TgPROP2 might not label mature autophagosomes. By analogy, in mammalian cells WIPI1 and WIPI2 were shown to be good autophagy markers as they are recruited to nascent autophagosomes upon amino acid deprivation, and WIPI-1 puncta-formation correlated with elevated levels of autophagosomal LC3 [31,32]; however, not all WIPI family members remain associated to mature autophagosomes [7,31].

**Fig 3.**
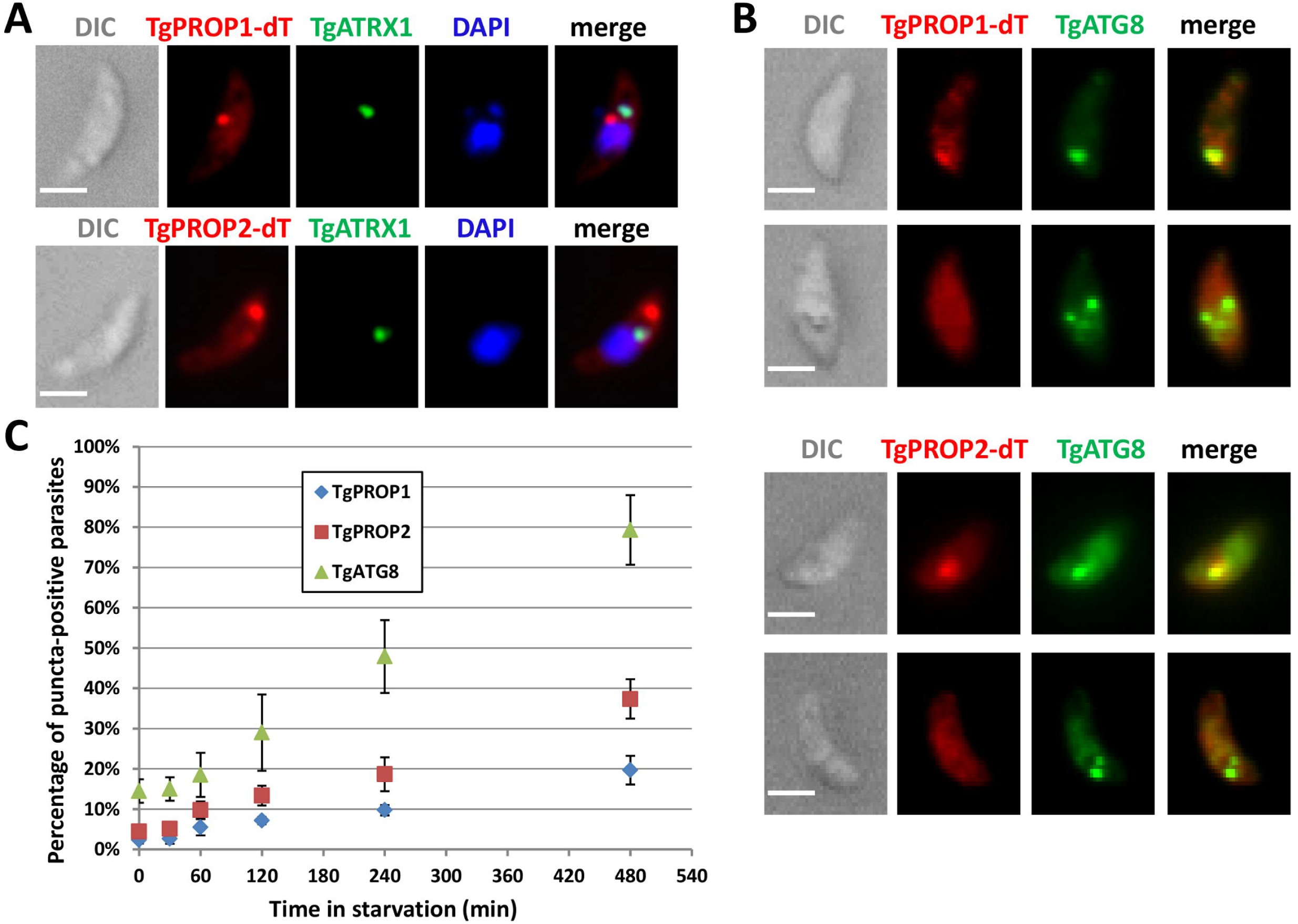
TgPROP1 and TgPROP2 relocalise to autophagic vesicles upon starvation. **A)** Extracellular parasites expressing dT-tagged TgPROP1 or TgPROP2 were incubated in amino acid-depleted medium for 6 hours before being fixed, adhered to poly-L-lysine slides, and processed for immunofluorescence using anti-TgATRX1 as an apicoplast marker. DNA was stained with DAPI. Scale bar= 2 μm. **B)** Extracellular parasites co-expressing GFP-TgATG8 together with dT-tagged TgPROP1 (top micrograph series) or TgPROP2 (bottom micrograph series), were incubated in amino acid-depleted medium for 6 hours before being fixed, adhered to poly-L-lysine slides, and processed for fluorescence imaging. DNA was stained with DAPI. Scale bar= 2 μm. **C)** Quantification of the percentage of extracellular parasites displaying a puncta-like signal for TgATG8, TgPROP1, or TgPROP2, during amino acid starvation for increased periods of time. Results are mean from *n*=3 experiments ±SEM.

### TgPROP1 and TgPROP2 vesicular localisation is PtdIns3P-dependent

Proteins of the Atg18/WIPI family are recruited to nascent autophagosomes through a conserved phosphoinositide-binding motif that preferentially binds to phosphoinositides such as PtdIns3P and, to a lesser extent, PtdIns(3,5)P_2_ [8,9,31,33]. Because the putative binding motif is conserved in TgPROP1 and TgPROP2, we expressed Td-fused mutated versions of these proteins where the two arginines in the binding motif were replaced by tyrosines. These mutated proteins remained essentially cytosolic when the parasites were placed in nutrient-depleted conditions (Fig 4A), suggesting their vesicular association is depending on phosphoinositides.

**Fig 4.**
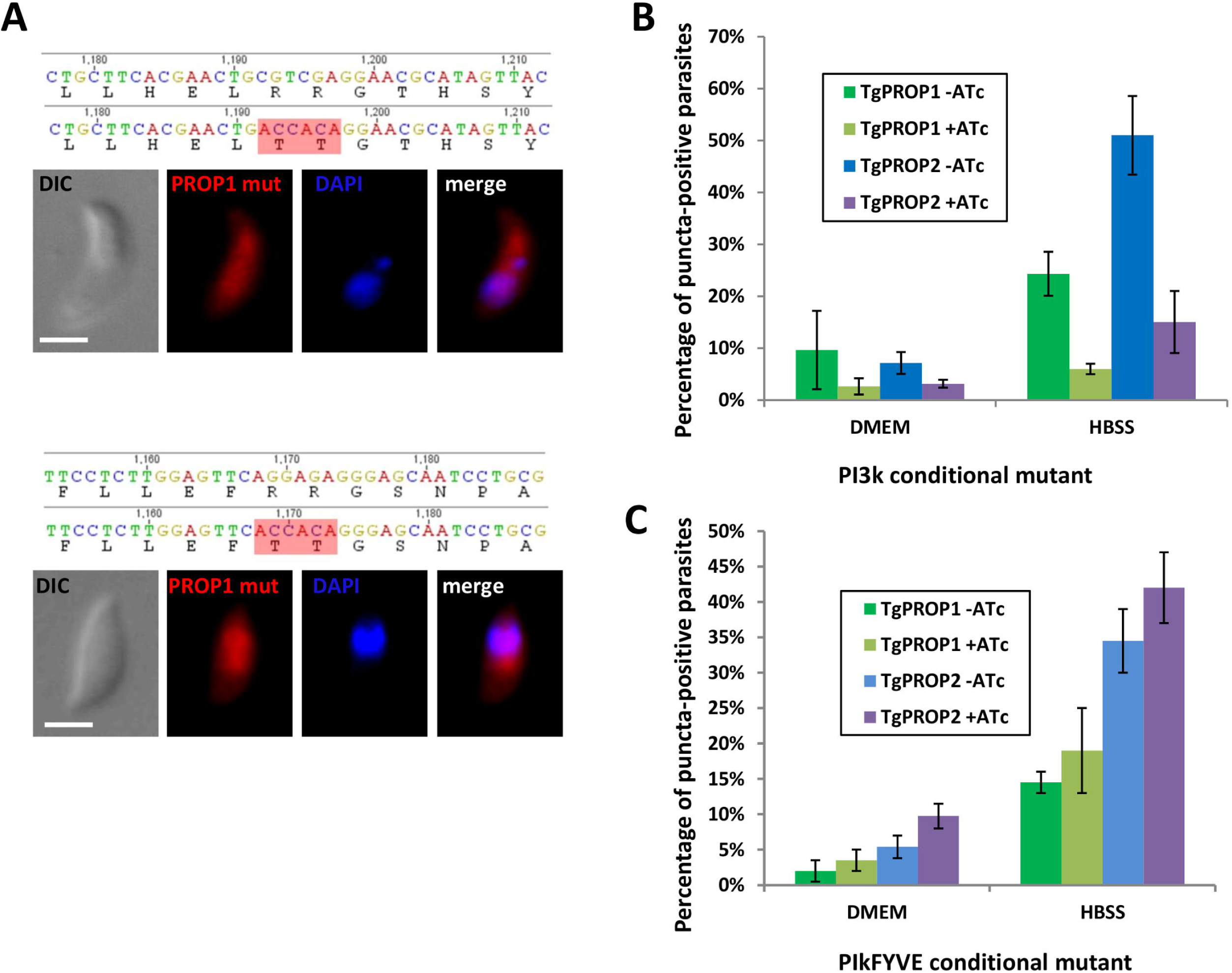
Binding of TgPROP1 and TgPROP2 to autophagic vesicles is PtdIns3P-dependent. **A)** dT-fused TgPROP1 (top) or TgPROP2 (bottom) mutated on the two arginine residues in their putative phosphoinositide binding motif do not localise anymore to vesicles in starved extracellular parasites. DNA was stained with DAPI. Scale bar= 2 μm. **B)** dT-fused TgPROP1 or TgPROP2 were expressed in the TgPI3k conditional mutant after depletion or not of the kinase by ATc; extracellular parasites were then starved as described before and the proportion of parasites with puncta-like TgPROP1 or TgPROP2 signal was evaluated. Results are mean from *n*=3 experiments ±SEM. **C)** dT-fused TgPROP1 or TgPROP2 were expressed in the TgPikFYVE conditional mutant after depletion or not of the kinase by ATc; extracellular parasites were then starved as described before and the proportion of parasites with puncta-like TgPROP1 or TgPROP2 signal was evaluated. Results are mean from *n*=3 experiments ±SEM.

To further verify his, we expressed dT-fused TgPROP1 and TgPROP2 in lipid kinases-deficient parasite cell lines. They were first expressed transiently in a class III phosphatidylinositol 3-kinase (P13k)- deficient cell line, which is unable to generate both PtdIns3P and PtdIns(3,5)P_2_ [34], put in nutrient depleted conditions, and puncta were quantified as described previously (Fig 4B). Depletion of TgPI3k activity severely impaired the formation of TgPROP1- and TgPROP2-positive puncta in these conditions (Fig 4B). PtdIns3P is also a precursor required for the production of PtdIns(3,5)P_2_ by a PI(3)P 5-kinase called PIKfyve in mammals, and a knock-down cell line for its homologue in *Toxoplasma* has also been generated [34], To decipher the relative importance of PtdIns3P or PtdIns(3,5)P_2_ for dT-fused TgPROP1 and TgPROP2 vesicular localisation, they were expressed in TgPikfyve-depleted parasites, which were put in nutrient depleted conditions, and puncta were quantified as described previously. In contrast with TgPI3k depletion, TgPikfyve depletion did not significantly affect TgPROP1- and TgPROP2-positive puncta formation (Fig 4B).

Overall, these data indicate vesicular localisation of TgPROP1- and TgPROP2 depends largely on their specific binding to PtdIns3P.

### Investigating TgPROP1 and TgPROP2 function

In order to get insights into the functions of TgPROP1 and TgPROP2, we sought to generate mutant cell lines. According to the recently published genome-wide CRISPR-based *Toxoplasma* phenotypic screen [35], TgPROP1 is likely to be a dispensable gene (positive phenotypic score), while TgPROP2 is probably important for parasite fitness *in vitro* and might be an essential gene (phenotypic score of - 1.46). We thus chose to generate inducible knock-down mutants in the TATi1-Ku80Δ background [36]. In a first step, to be able to check for efficient protein depletion, we added a sequence coding for a C-terminal triple hemagglutinin (HA) epitope tag at the endogenous *TgPROP1* or *TgPROP2* loci by single homologous recombination (Fig 5A) to yield the TgPROP1-HA and TgPROP2-HA cell lines, respectively. Transfected cKd-TgPROP1-HA and cKd-TgPROP2-HA parasites were subjected to chloramphenicol selection and clones were isolated and checked for correct integration of the plasmid by PCR (Fig 5B). Then, in these cell lines we replaced the endogenous promoter by an inducible-Tet07*SAG*4 promoter, through a single homologous recombination at the *TgPROP1* or *TgPROP2* loci, to yield conditional TgPROP1 and TgPROP2 knock-down cell lines (cKd-TgPROP1-HA and cKD-TgPROP2-HA, respectively) (Fig 5A). The addition of anhydrotetracycline (ATc) would then repress *TgPROP* transcription through inhibition of the TATi1 transactivator and subsequently block the Tet-operator [37], Several pyrimethamine-isolated clones were selected after diagnostic PCR for correct genomic integration (Fig 5C).

**Fig 5.**
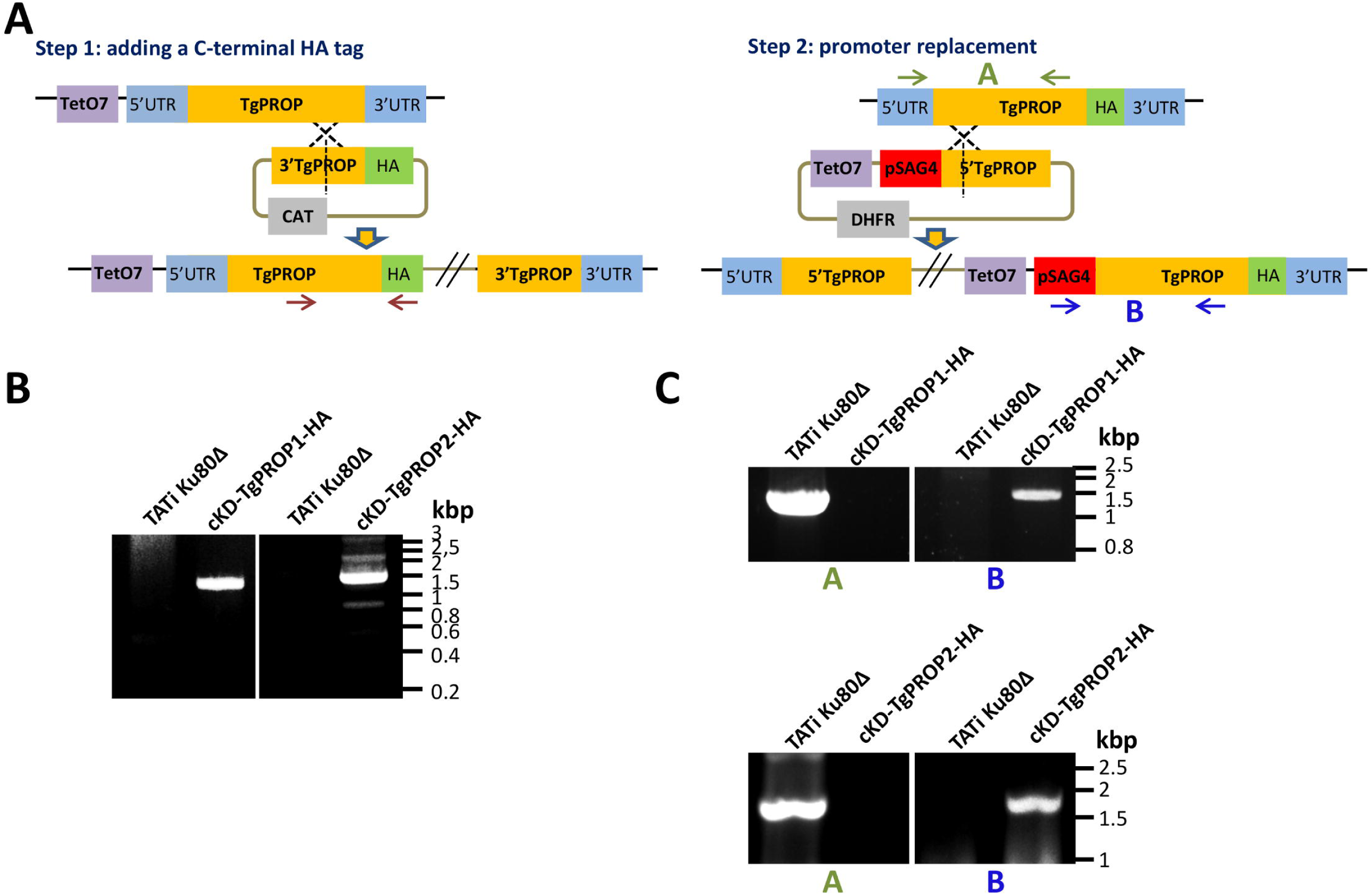
Generation of HA-tagged conditional mutant cell lines for TgPROP1 and TgPROP2. **A)**Schematic representation of the two-steps general strategy for C-terminal HA tagging of TgPROP1 and TgPROP2 and subsequent generation of conditional mutant by replacement of the native promoter by an ATc-regulatable one. **B)** Diagnostic PCR for verifying proper integration of the construct for the C-terminal HA-tagging of TgPROP1 and TgPROP2. **C)** Diagnostic PCRs for *TgPROP1* (top) or *TgPROP2* (bottom) promoter replacement.

### TgPROP1 seems involved in parasite stress response, but is not essential for *in vitro* growth

Anti-HA antibodies were used to detect tagged TgPROP1 by immunoblot, and revealed a product larger than its expected molecular mass (100 kDa versus 72 kDa, Fig 6A). It should also be noted that the antibodies detected a second, smaller product of about 40 kDa. We were wondering if this could be due to the promoter change, leading potentially to a different timing or level of expression during the cell cycle that would cause an unusual processing or degradation of the main product. To verify this, we checked for protein expression in the TgPROP1-HA cell line, where the tagged protein is under the control of its own promoter. Immunoblot analysis revealed that the main product corresponding to TgPROP1-HA is indeed of about 100 kDa, and that the promoter change likely led to the appearance of the smaller product, as the latter was essentially absent when the tagged protein was expressed from its native promoter (S1A Fig). This also allowed us to verify that both under its native promoter, or after the *SAG4* promoter exchange, TgPROP1-HA localises in the cytoplasm of intracellular parasites, and relocalises to puncta in extracellular parasites after amino acid depletion (S1B-C Fig), as observed with the dT-fused protein. Immunoblot and immunofluorescence analyses showed an efficient depletion of both TgPROP1 upon ATc treatment for two days (Fig 6A-B).

**Fig 6.**
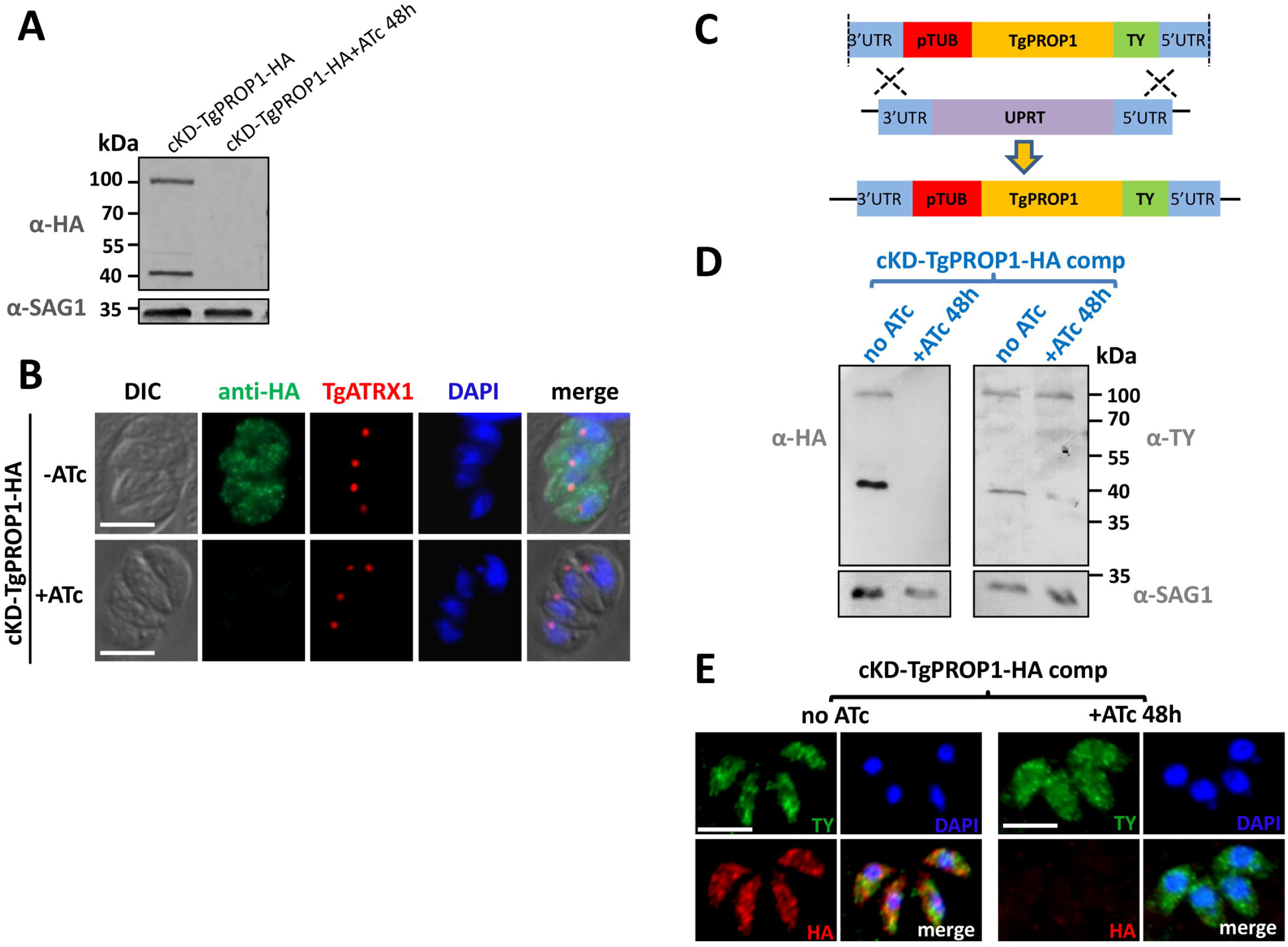
Regulation of TgPROP1 and TgPROP2 depletion. **A)** Immunoblot verification of TgPROP1 depletion after two days of ATc treatment. **B)** Verification by fluorescence microscopy of efficient TgPROP1 depletion after two days of ATc treatment. TgATRX1 was used as a marker to check for apicoplast integrity. DNA was stained with DAPI. Scale bar= 5 μm. **C)** Schematic representation of the strategy for complementing TgPROP1 conditional mutant by integrating a copy at the *UPRT* locus for expressing a TY-tagged version of the protein. **D)** Immunoblot showing the TY-tagged TgPROP1 copy is expressed at the expected molecular mass and remains expressed upon depletion of the regulatable HA-tagged copy by two days of ATc treatment. SAG1 was used as a loading control. **E)** Specific immunodetection of TY-tagged TgPROP1 (green) in intracellular parasites upon depletion of the regulated HA-tagged copy (red) after two days of ATc treatment. DNA was stained with DAPI. Scale bar= 5 μm.

We also generated a complemented cell line expressing constitutively an additional copy of *TgPROP1* from the *uracil phosphoribosyltransferase* (*UPRT*) locus (Fig 6C), in the conditional mutant background. After depletion of the ATc-regulated copy, this cell line maintained expression of a TY-tagged TgPROP1 with its expected molecular mass and localisation (Fig 6D-E).

In contrast with the apicoplast-related TgATG mutants previously characterised [18,22,23], ATc-driven depletion of TgPROP1 did not affect the parasite lytic cycle in vitro (Fig 7A). Consistent with this, we found that conditional depletion of TgPROP1 had no particular impact on the apicoplast (Fig 6B). This was similar to what we previously observed with the *Toxoplasma* homologue of ATG9, another early component of the autophagy machinery [24], TgATG9 was nevertheless shown to be important for the parasite in particular stress conditions or in the context of the host [24], so we next investigated if that could also be the case for TgPROP1. The viability of the cKd-TgPROP1-HA cell line was significantly reduced after prolonged exposure to an extracellular stress (Fig 7B). There was already some reduction in viability upon stress before depleting the protein with ATc, which we think might be explained by the reduced expression levels and the potential protein cleavage that occur after promoter change (S1 Fig). When we performed invasions with freshly egressed parasites, cKd-TgPROP1-HA parasites behaved as the parental cell line (S2 Fig), demonstrating viability of the cKd-TgPROP1-HA cell line is specifically affected by a prolonged extracellular state.

**Fig 7.**
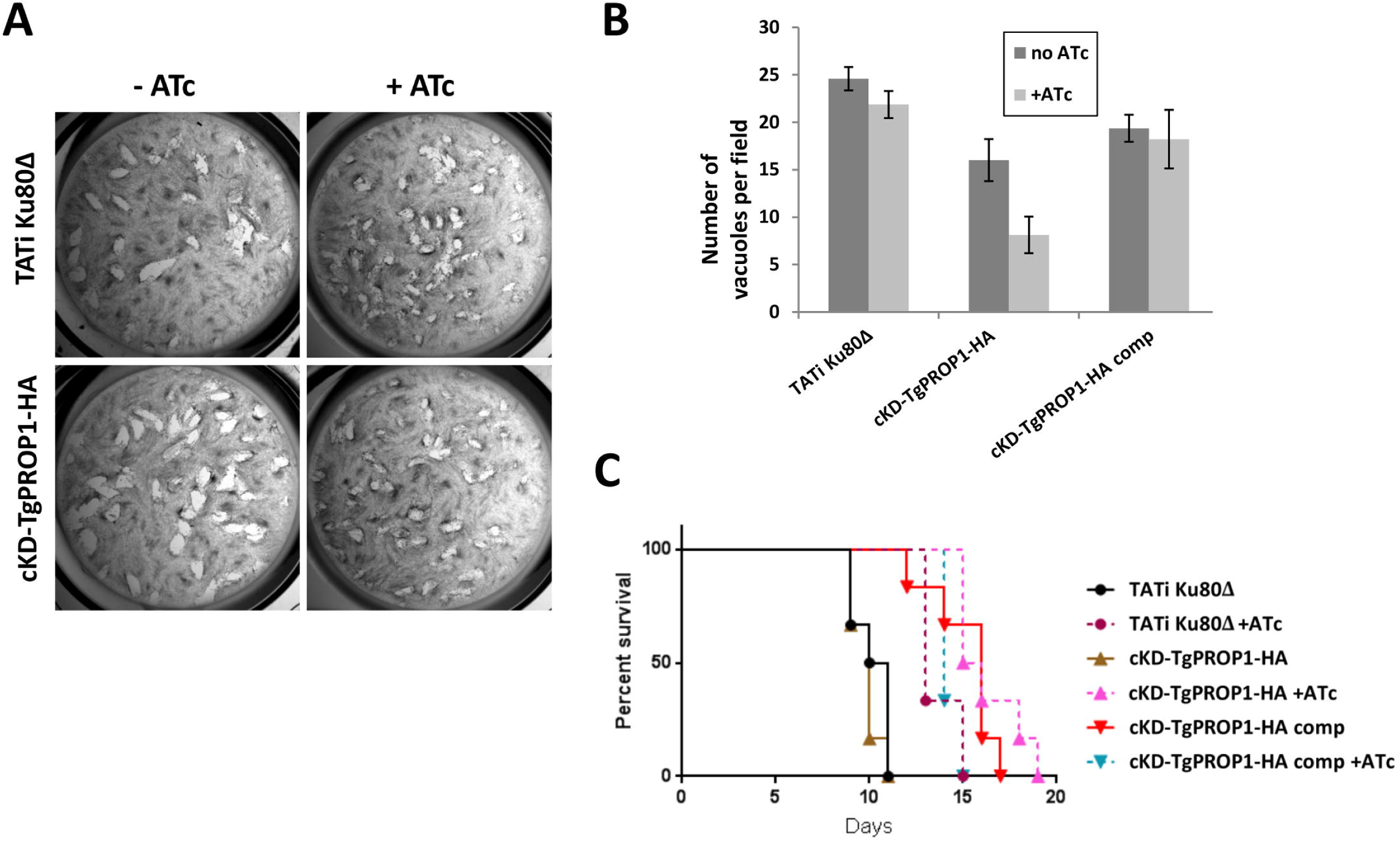
Phenotypic analysis of TgPROP1-depleted parasites shows the protein could be involved in the stress response. **A)** Plaque assays were carried out by infecting HFF monolayers with parasites from the conditional mutant cell lines kept or not in the presence of ATc for the duration of the experiment. After 6 days, plaques are visible in the host cells monolayer, suggesting an unimpaired lytic cycle. **B)** Viability of parasites after 16 hours of extracellular incubation in DMEM was assessed by their ability to invade host cells. TATi-Ku80Δ and cKD-TgPROP1-HA parasites were kept or not with ATc for two days. Data shown are the mean values ± SEM from one representative experiment out of three. **C)** Mouse infectivity assay. Survival curve of BALB/c mice (*n*=**5)** infected intraperitoneally with 1000 parasites from the TgPROP1 knock-down and complemented cell lines kept, or not, in the presence of ATc. Data are representative of three independent experiments.

We next tested if depletion of TgPROP1 could have an impact on parasite virulence in the mouse model (Fig 7C). All mice died after being injected with the various cell lines. However, depletion of cKd-TgPROP1-HA led to a slight delay in death that was consistently observed in the three independent experiments we performed.

Overall, TgPROP1 do not seem to be essential for normal parasite growth *in vitro* and survival *in vivo*, although depletion of TgPROP1 renders the parasites less able to cope with extracellular stress and might reduce their fitness in mice.

### TgPROP2 is potentially important for parasite viability *in vitro*

Upon HA-tagging, detection by immunoblot TgPROP2 revealed a product of 70 kDa, slightly larger than the theoretical molar mass of 60 kDa (S3A Fig and S3D Fig), similar to what we observed for TgPROP1-HA (Fig 6A) and the dT-fused versions of *Toxoplasma* PROPPINs (Fig 2A). We then performed a phenotypic analysis for the conditional mutant cell line we generated for TgPROP2. The protein was apparently efficiently depleted upon addition of ATc (S3A-B Fig and S3D-E Fig), yet like for TgPROP1 there was no apparent *in vitro* growth phenotype as assessed by plaque assay (S3C Fig). This was somewhat unexpected, as a CRISPR screen predicted a potential fitness cost after abolishing the expression of this protein [35]. We thus conducted additional immunoblot, immunofluorescence and RT-PCR analyses (S3D-F Fig) and noticed that the promoter replacement in the conditional knockdown cell line led to a significant increase in mRNA and protein expression, which was efficiently subsequently down-regulated by the addition of ATc to levels beyond detection by immunoblot or immunofluorescence. However, because of the presence of residual amounts of mRNA after several days of ATc treatment (S3F Fig), we could not rule out the possibility that minute amounts of TgPROP2 were still expressed and functional in the knock-down parasites. Moreover, in contrast with *TgPROP1* (Fig S4), and in spite of multiple attempts, we failed to generate a stable *TgPROP2* knockout cell line by CRISPR-driven recombination, which highlighted a potential essential role for the corresponding protein. Besides, when checking for the presence of the apicoplast 48 hours after transfection, about 30% of the vacuoles had parasites lacking the organelle. This is in accordance with a recently published study describing an essential function for TgPROP2 (simply called ‘TgATG18’ in the aforementioned study) for apicoplast maintenance and parasite viability [38].

## DISCUSSION

Autophagy is one of the most conserved cellular pathways in eukaryotes. However, in spite of accumulating evidence suggesting this pathway is likely functional and would be part of an integrated stress response in *Toxoplasma,* the clear demonstration of its catabolic function has not been made in Apicomplexa. The autophagy-related molecular machinery is very peculiar in these parasites. First, they seem to have only partly retained the canonical machinery initially described in yeast and other model organisms [17]. Second, several Apicomplexa have been repurposing part of this machinery for an autophagy-independent function [21,23,39].

For instance, our previous research has shown that TgATG8 associates with the outermost membrane of the apicoplast [21]. Moreover this protein, along with other proteins regulating its membrane association, are crucial for maintaining the homeostasis of this important organelle and thus for viability of the parasites [21,22], In addition, TgATG8 localises to autophagic vesicles upon nutrient starvation [18], which suggests it is also involved in canonical autophagy. However, in contrast to the ATG8-centered machinery involved in an apicoplast-related function, early autophagy marker TgATG9 was found to be not essential for *in vitro* growth, albeit potentially important for surviving extracellular stresses and for fitness in mice [24], This suggested canonical autophagy is not essential to parasite growth and prompted us to study the function of other proteins involved in early stages of autophagosome formation, like lipid sensors of the PROPPIN family, that are important scaffold proteins for the expansion of autophagic vesicles.

Here we show that TgPROP1 and TgPROP2 are relocalising to vesicular structures upon starvation, and partially co-localise with TgATG8, which suggest these vesicles could be of autophagosomal nature. We also demonstrated this membrane association is dependent on the lipid PtdIns3P. Contrarily to TgATG8, however, these TgPROPs are not found at the apicoplast. We demonstrated here that depletion of TgPROP1 has little impact of parasite viability *in vitro* in normal growth conditions. Interestingly, however, TgPROP1-depleted parasites seem to be less able to cope with stress as extracellular parasites, and might be slightly less fit in the animal host. This is somewhat reminiscent of the TgATG9 mutant, although the latter has a more pronounced virulence phenotype in mice [24], Altogether, this would indicate that, like TgATG9, TgPROP1 is primarily involved in the canonical autophagy pathway, which is not essential for growth in nutrient-rich culture conditions in most eukaryotic cells.

While TgPROP2 can be recruited to autophagic vesicles like TgPROP1, in sharp contrast with the latter it seems to be more important for parasite growth *in vitro.* The conditional knock-down cell line we generated had no obvious growth phenotype *in vitro* (S3 Fig), however we failed at isolating*TgPROP2* knock-out parasites after multiple attempts, while a *TgPROP1* knock-out cell line was easily obtained (S4 Fig). Moreover, a CRISPR-based *Toxoplasma* phenotypic screen also suggests it is important for fitness [35]. Besides, an independent functional analysis of the single PROPPIN homologue of *Plasmodium*, and of TgPROP2 (called PfATG18 and TgATG18, respectively, by their authors) was recently published [38], describing an important function for these proteins in the maintenance of the apicoplast and the viability of the parasites. Altogether this points towards an important function for TgPROP2. However it seems only minute amounts of protein might be necessary for its function: in our conditional knock-down cell line, although mRNA was still present, the protein was not detectable by immunological methods, yet the parasites were viable.

Bansal et al. suggested TgATG18 has a potential influence on TgATG8 lipidation and its association with the apicoplast [38]. In other eukaryotes, several studies have also suggested PROPPINs can regulate Atg8/LC3 lipidation at the autophagosomal membrane [40,41], but they can be found together at the nascent organelle. At this stage it is unclear how a PROPPIN that does not localise to the apicoplast would regulate TgATG8 conjugation at the membrane of the organelle. The lipid-biding properties of PROPPINs are thought to be key for exerting their function [40,41], We demonstrated here that the lipid-binding motif of TgPROPs, and TgPI3k (but not TgPIKfyve) activity were important for vesicular localisation of these proteins. Bansal et al. also found that recombinant TgATG18/TgPROP2 is able to bind PtdIns3P through the lipid binding motif present in the WD-40 domain of the protein, and showed this is important for the function they describe in apicoplast homeostasis [38]. Interestingly, PtdIns3P has been known for some time to be a critical player in maintaining apicoplast homeostasis [28]. We have previously shown through functional analyses of lipid kinase mutants that the downstream lipid effector PtdIns(3,5)P_2_ is important for apicoplast integrity, although a direct role of PtdIns3P in recruiting apicoplast effectors cannot be excluded [34], How phosphoinositides would regulate an apicoplast-related function for *Toxoplasma* PROPPINs is, however, not clearly established at the moment. Several known TgATG8 effectors (TgATG4, TgATG3) have not been observed at the apicoplast [18,22], and TgATG8 recruitment at the membrane of the organelle seems to be regulated during the cell cycle [21], It is thus possible several modulators of TgATG8 association with the apicoplast membrane are acting transiently or in low amounts, and are difficult to detect at the organelle.

Homologous proteins of the PROPPIN family can be found in several distinct clades of the eukaryotic kingdom, which suggests this family is ancient. Several studies have classified yeast and human PROPPINs in two paralogous groups: one containing human WIPI1, WIPI2 and the ancestral yeast Atg18, and the other containing human WIPI3 and WIPI4 [10,33]. Although proteins from these two groups sometimes have a relatively limited homology outside of their WD-40 domain, their role generally gravitates around several different autophagy-related processes. Their functions have remained incompletely understood at the molecular level and recent investigations suggest PROPPINs could be involved in cellular mechanisms as diverse as membrane fission [42], or scaffolding for signaling cascades [43]. It is interesting to note that in *Plasmodium* there is a single member of the PROPPIN family, while *Toxoplasma* has two, of which only one (TgPROP2) might be involved in an apicoplast-related function. TgPROP1, on the other hand, might be exclusively involved in a more canonical autophagy pathway. Interestingly, TgPROP1 appears to be the closest phylogenetic relative of yeast Atg18 and mammalian WIPI2 (Fig 1C), which are the prototypical autophagy-related PROPPINs. Whether or not the *Plasmodium* PROPPIN is also involved in the biogenesis of autophagosomes besides its apicoplast-related function is unknown, and this raises the possibility of a specific functional diversification in selected members of the phylum Apicomplexa.

Whether it is to elucidate the apicoplast-related function of some of the autophagy-related proteins, or to establish more firmly the presence of a canonical autophagy pathway, further studies are necessary to dissect functionally this molecular machinery in apicomplexan parasites. However, the example of the PROPPIN family members illustrates how tedious this task is: as some members of this machinery might be involved exclusively in the autophagy pathway, while others are likely to have moonlighting activities and thus are also involved in distinct, yet intertwined, cellular functions.

## MATERIALS AND METHODS

### Ethics statement

All murine experiments were approved by the Laboratory Animal Ethics Committee of Wenzhou Medical University (Permit number #wydw 2016-118). Mice were housed in strict accordance with the Good Animal Practice requirements of the Animal Ethics Procedures and Guidelines of China. Humane endpoints were used to avoid the mice pain or suffering via euthanasia. Mice were monitored twice each day for signs of toxoplasmosis, such as impaired mobility, difficulty in feeding, weight loss, self-mutilation and severe ascites. Mice were sacrificed immediately with CO2 gas when shown above signs. Eye pricks were performed following deep anesthesia.

### Parasites and cells culture

Tachyzoites of the RH HxGPRTΔ [44], TATi1-Ku80Δ [36], pi3ki or pikfyvei [34] *T. gondii* cell lines, as well as derived transgenic parasites generated in this study were maintained by serial passage in human foreskin fibroblast (HFF, American Type Culture Collection, CRL 1634) cell monolayer grown in Dulbecco’s Modified Eagle’s medium (DMEM, Gibco) supplemented with 5% decomplemented fetal bovine serum (Gibco), 2mM L-glutamine (Gibco) and a cocktail of penicillin–streptomycin (Gibco) at 100 μ/mL.

### Bioinformatic analyses

Sequence alignments were performed using the multiple sequence comparison by log-expectation algorithm of the Geneious software suite v5.3.3 (http://www.geneious.com). The phylogenetic tree was built with the same software suite using Jukes-Cantor genetic distance model, and unweighted pair group method with arithmetic mean for tree building. Domain searches were performed in the Pfam database (http://pfam.xfam.org/).

### Generation of Tomato dimer-tagged TgPROP1 and TgPROP2 cell lines

Plasmids TgPROP1-dT and TgPROP2-dT, for expressing TgPROP1 and TgPROP2, respectively, fused with a C-terminal dimeric Tomato fluorescent protein, under the dependence of a *tubulin* promoter were constructed as follows. TGGT1_288600 (TgATG8a, 1968 bp) or TGGT1_220160 (TgPROP2, 1662 bp) open reading frames were amplified by PCR from cDNA using primers ML1671/ML1672 or ML1643/ML1644 (see S1 Table for primer sequences), respectively, and cloned into the BgIII and Avrll sites in the pTub-IMC1-dimeric Tomato.CAT vector [21]. The RH HxGPRTΔ [44], pi3ki or pikfyvei [34] cell lines were transfected with 100 μg of each plasmid, and then subjected to chloramphenicol selection. Site-directed mutagenesis was performed with the Quikchange mutagenesis kit (Agilent) using primers ML2293/ML2294 and ML2295/2296, to introduce mutations in the TgPROP1-dT and TgPROP2-dT, respectively, in order to express versions of the PROPPINs mutated in their lipid binding site.

### Generation of conditional knock-down, complemented and HA-tagged TgPROP1 and TgPROP2 cell lines

TgPROP1 and TgPROP2 were tagged in their respective conditional mutant background. The 3’ end of *TgPROP1* was amplified by PCR from genomic DNA, with the Q5 polymerase (New England Biolabs) using primers ML2488/ML2489 and inserted in frame with the sequence coding for a triple HA tag present in the pLIC-HA_3_-CAT plasmid. The resulting vector was linearized with SphI, and 50 μg of DNA were transfected into the cKD-TgPROP1 cell line. Parasites were selected by chloramphenicol to yield the cKD-TgPROP1-HA cell line. The 3’ end of *TgPROP2* was amplified by PCR from genomic DNA, with the Q5 polymerase (New England Biolabs) using primers ML1111-ML1112 and inserted in frame with the sequence coding for a triple HA tag, present in the pLIC-HA_3_-CAT plasmid. The resulting vector was linearized with MluI and 50 μg of DNA were transfected into the cKD-TgPROP2 cell line. Parasites were selected by chloramphenicol to yield the cKD-TgPROP2-HA cell line.

To generate the cKD-TgPROP1 and cKD-TgPROP2 conditional mutant parasites, two genomic fragments of 1039 and 1631 bp corresponding to the 5’ genomic sequences of the *TgPROP1* or *TgPROP2* genes, respectively, were amplified by PCR using primers ML1399/ML1400 or ML1496/ML1497 and subcloned into BglII and NotI sites of ptet07Sag4-HA(2) vector [36]. The resulting plasmids were named. TATi1-Ku80Δ parasites were transfected with 30 μg of each vector (linearized with AatII and NsiI respectively) and subjected to pyrimethamine selection. Correct integration at the locus was verified by PCR using primers ML1041/ML1445 (for cKD-TgPROP1) and primers ML1041/ML3110 (for cKD-TgPROP2). Control amplification of the wild-type 5’ region was performed with primers ML1444/ML1445 (for TgPROP1) and ML3109/ML3110 (for TgPROP2).

The cKD-TgPROP1-HA cell line was complemented by the addition of an extra copy of *TgPROP1* at the *uracil phosphoribosyltransferase* (*UPRT)* locus. The TgPROP1-dT plasmid generated previously was used to obtain the *TgPROP1* sequence by AvrII/BgIII digestion, and the fragment was cloned into the Avrll/Bglll-digested pUPRT-TUB-Ty vector [36]. This plasmid was then digested to provide a donor sequence to be integrated by CRISPR/CAS9. To do this, this was co-transfected with the pSS013-CAS9NLS-YFP plasmid (kind gift from B. Striepen and M. Cipriano, University of Georgia, USA), expressing a *UPRT*-specific guide RNA under the control of a U6 promoter, and expresses a nuclear localized Cas9-YFP. Transgenic parasites were selected using 5-fluorodeoxyuridine.

### Inactivation of *TgPROP1* and *TgPROP2* by CRISPR-mediated direct knock-in strategy

Guide sequences were designed to target the 5’ of *TgPROP1* using primer couple ML2766/2767, or both the 5’ and 3’ of *TgPROP2* with primer couples ML3223/ML3224 and ML3225/ML3226, respectively. Guide sequences were cloned into the pU6-universal plasmid (kind gift from S. Lourido, Whitehead Institute, USA). For *TgPROP1,* the donor sequence was obtained by PCR with the KOD DNA polymerase (Merck), using primers ML2770/ML2771, with the pSAG1-DHFR plasmid as a template. For *TgPROP2,* the donor sequence was obtained by PCR with the KOD DNA polymerase (Merck), using primers ML3350/ML3352, with plasmid pT8MycGFP-HX [45] (kind gift of D. Soldati-Favre, University of Geneva, Switzerland) as a template. Approximately 40 μg of each guide and 5 μg of donor DNA were co-transfected.

### Immunoblot analysis

Proteins from freshly lysed parasites (10^7^ per sample) were separated by SDS-PAGE and analysed by immunoblot as described previously [46]. Epitope-tagged TgPROP1 and TgPROP2 were detected with rat monoclonal anti-HA (Roche), rabbit polyclonal anti-RFP (Abcam), or mouse anti-Ty [47] antibodies. Mouse anti-SAG1 monoclonal antibody [48] or anti-TgIF2α [49], were used to detect the loading control.

### Plaque assays

Confluent monolayers of HFFs grown in 24-well plates were infected with 2 × 10^5^ freshly egressed tachyzoites and incubated with or without ATc (at 1.5 μg/ml) for 6 days. Infected cell layer was then fixed in cold methanol and stained with Giemsa. Images were acquired with an Olympus MVX10 macro zoom microscope equipped with an Olympus XC50 camera.

### Parasite viability assays

Freshly lysed tachyzoites of TATi-Ku80Δ, cKD-TgPROP1 and cKD-TgPROP2 cell lines were were incubated in complete DMEM or amino acid depleted Hank’s Balanced Salt Solution (HBSS) at 37 °C with 5% C0_2_ for 16 hrs. For assessing parasite viability, we then evaluated their invasive capacity as described previously [24], They were counted and 2 × 10^5^ were used to infect confluent monolayer of HFFs grown on coverslips for 18 hrs. The number of parasitophorous vacuoles per field was visualized by immunofluorescence assay (IFA) using anti-ROP1 antibody, with a 100× objective lens.

### Immunofluorescence microscopy

For IFA, intracellular tachyzoites grown on coverslips containing HFF monolayers were fixed for 20 min with 4% (w/v) paraformaldehyde in PBS, permeabilised for 10 min with 0.3% Triton X-100 in PBS and blocked with 0.1% (w/v) BSA in PBS. Primary antibodies used for detection of the the apicoplast were mouse monoclonal anti-ATRX1 (1/1000, kind gift of Peter Bradley, UCLA, USA) and rabbit anti-TgCPN60 (1/2000) [50]. Rat monoclonal anti-HA antibody (clone 3F10, Roche) was used at 1/500 to detect HA-tagged TgPROP1 and TgPROP2 and mouse anti-Ty [47] to detect Ty-tagged TgPROP1. Staining of DNA was performed on fixed cells incubated for 5 min in a 1 μg/ml DAPI solution. All images were acquired at the “Montpellier Ressources imagerie” facility from a Zeiss AXIO Imager Z2 epifluorescence microscope equipped with a Camera ORCA-flash 4.0 camera (Hammamatsu) and driven by the ZEN software. Adjustments for brightness and contrast were applied uniformly on the entire image.

### Analysis of autophagosome-like structures

To visualize autophagosome-like structures, *T. gondii* tachyzoites were transfected to express GFP-TgATG8 [18] and costained with antibodies for the TgATRX1 apicoplast marker [30] to discard the apicoplast-related signal.

### Semi-quantitative RT-PCR

RNAs were extracted from extracellular *T. gondii* tachyzoites, incubated with or without ATc for 3 days, using the Nucleospin RNA II Kit (Macherey-Nagel). cDNAs were produced with 800 ng of total RNA per RT-PCR reaction using the Superscript II first-strand synthesis kit (Invitrogen). Specific primers for *TgPROP2* (ML3245/ML3246), *TgPROP1* (ML3270/ML3271) and, as a control, β*-tubulin* (ML841/ML842) primers, were used for PCR with the GoTaq polymerase (Promega). 23 cycles of denaturation (10 s, 95°C), annealing (30 s, 55°C) and elongation (15 s, 68°C) were performed.

### Virulence assays in mice

Survival experiments were performed on groups of 6 mice per parasite cell line. Six to eight weeks old female C57BL/6 mice, purchased from Laboratory Animal Center of Wenzhou Medical University, were infected by intraperitoneal (i.p.) injection of 1 000 tachyzoites freshly harvested from cell culture. ATc (Cayman CAS: 13803-65-1) was added to the drinking water at a final concentration of 0.2 mg/ml and the solution changed every 36 hrs. The water bottle containing ATc was wrapped in aluminum foil to prevent precipitation of ATc due to light. Mice survival was checked twice daily until their death, endpoint of all experiments. On day 10 postinfection, sera from surviving mice was monitored for immune response by immunoblotting against tachyzoites lysates. Data were represented as Kaplan and Meier plots using GraphPad Prism 6.0.

## ACKNOWLEDGEMENTS

We wish to thank B. Striepen and M. Cipriano, S. Lourido, D. Soldati-Favre, L. Sheiner, V. Carruthers, JF Dubremetz, W. Sullivan and P. Bradley for their kind gift of cell lines, plasmids or antibodies. We also thank the “Montpellier Ressources imagerie” platform for providing access to their microscopes.

## SUPPORTING INFORMATION CAPTIONS

**S1 Fig. Tagging of TgPROP1 at the endogenous locus. A)** Immunoblot analysis of the HA-tagged TgPROP1 expressed with its own promoter (TgPROP1-HA) or after replacement with a *SAG4* promoter (cKD-TgPROP1-HA). SAG1 was used as a loading control. **B)** Localisation of HA-tagged TgPROP1 expressed with its own promoter (TgPROP1-HA) or after replacement with a *SAG4* promoter (cKD-TgPROP1-HA) in intracellular (left) or starved extracellular (right) parasites. DNA was stained with DAPI. Scale bar = **5** μm (left) or 2 μm (right).

**S2 Fig. Freshly egressed cKD-TgPROP1-HA parasites retain full invasive capacity.** TATi-Ku80Δ, cKD-TgPROP1-HA and cKD-TgPROP2-HA parasites were mechanically released from their host cells and assessed for their ability to invade host cells.

**S3 Fig. Conditional cKD-TgPROP2-HA cell line has no obvious** *in vitro* **growth phenotype.**

**(A)**Immunoblot analysis of TgPROP2-HA depletion after two days of ATc incubation. SAG1 was used as a loading control. **B)** IFA of TgPROP2-HA depletion after two days of ATc incubation. TgPROP2 was detected with anti-HA antibodies, the apicoplast was detected using anti-TgATRX1 antibodies. DNA was stained with DAPI. Scale bar= 5 μm. **C)** Plaque assay show conditional depletion of TgPROP2 has no drastic effect on the lytic cycle. **D)** Immunoblot analysis shows promoter change leads to an overexpression of TgPROP2-HA. TgIF2α was used as a loading control. **E)** IFA also shows a higher level of TgPROP2-HA expression when expressed from the *SAG4* promoter. DNA was stained with DAPI. Scale bar= 5 μm. **F)** Semiquantitative RT-PCR analysis of *TgPROP2* expression shows minute amounts of mRNA are still detectable after 3 days of incubation with ATc. Analysis was performed on parasites incubated or not with ATc for 3 days regulate mRNA expression. Specific β*-tubulin* primers were used as controls.

**S4 Fig. Generation of a** *TgPROP1* **knock-out mutant. A)** Schematic representation of the strategy for generating a *TgPROP1* knock-out cell line using CRISPR/Cas9. Locus modification was made in the RH cell line deleted for the *Hypoxanthine-guanine phosphoribosyltransferase* (*HxGPRT*) gene. The donor sequence contained a HxGPRT sequence for selection of transgenic parasites with mycophenolic acid and xanthine. **B)** RT-PCR analysis showing efficient *TgPROP1* mRNA depletion in the *TgPROP1Δ* cell line. Specific *TgPROP2* and β*-tubulin* primers were used as controls.

**S1 Table. Primers used in this study.**

